# A Single Cell Atlas of Lung Development

**DOI:** 10.1101/2021.01.21.427641

**Authors:** Nicholas M. Negretti, Erin J. Plosa, John T. Benjamin, Bryce A. Schuler, A. Christian Habermann, Christopher Jetter, Peter Gulleman, Chase J. Taylor, David Nichols, Brittany K. Matlock, Susan H. Guttentag, Timothy S. Blackwell, Nicholas E. Banovich, Jonathan A. Kropski, Jennifer M. S. Sucre

## Abstract

Lung organogenesis requires precisely timed shifts in the spatial organization and function of parenchymal cells, especially during the later stages of lung development. To investigate the mechanisms governing lung parenchymal dynamics during development, we performed a single cell RNA sequencing (scRNA-seq) time-series yielding 92,238 epithelial, endothelial, and mesenchymal cells across 8 time points from embryonic day 12 (E12) to postnatal day 14 (P14) in mice. We combined new computational analyses with RNA *in situ* hybridization to explore transcriptional velocity, fate likelihood prediction, and spatiotemporal localization of cell populations during the transition between the saccular and alveolar stages. We interrogated this atlas to illustrate the complexity of type 1 pneumocyte function during the saccular and alveolar stages, and we demonstrate an integrated view of the cellular dynamics during lung development.

## Introduction

Organogenesis of the lung requires precisely timed and coordinated signaling between multiple cell types, as well as the adaptation of cellular functions from intrauterine to extrauterine life after birth (Morrisey and Hogan, 2010). The later aspects of lung development are exquisitely fluid, with dramatic changes in spatial organization and function of epithelial, endothelial, and mesenchymal cells that coordinate sacculation and alveolarization (Warburton et al., 2010). Despite considerable progress, an integrated understanding of the identity, localization, and fate of alveolar parenchymal cells during lung development has remained elusive.

During the late terminal saccular stage (24 weeks to later fetal period in humans; embryonic day (E)17.5 to postnatal day (P) 5 in mice) (Warburton et al., 2010), the lung is especially vulnerable to injury. The developmental program of the lung in the preterm infant can be permanently altered with exposure to hyperoxia and inflammation, resulting in persistent structural damage and lifelong respiratory impairment (Benjamin et al., 2020; Goss, 2018). Presently, the molecular mechanisms that alter the normal developmental trajectory are not well understood. There is a critical need to elucidate the complex interactions that occur between cellular sub-populations in order to provide context for understanding the functional mechanisms regulating lung development, homeostasis, injury, and pathology.

To develop a more comprehensive and integrated understanding of the changes in cellular dynamics during lung development, we performed single cell RNA sequencing (scRNA-seq) on 92,238 cells from lung tissue across 8 developmental timepoints from E12 to P14. By applying transcriptional velocity analysis (La Manno et al., 2018) together with fate likelihood prediction (Lange M, 2020) and RNA-in situ hybridization, we characterize the transcriptomic trajectories of epithelial, endothelial, and mesenchymal cells across developmental time. We identify the emergence of a transitional population of epithelial cells beginning at E15, suggesting early alveolar type 1 epithelial (AT1) cell fate commitment, illustrate the developmental trajectory of the recently described *Car4*+ endothelial cells, and demonstrate the transitions in Wnt ligand expression that accompany population shifts in the pulmonary mesenchyme during alveologenesis. We interrogate this cellular atlas to describe a previously unanticipated role of alveolar type 1 pneumocytes (AT1) in the expression of genes associated with elastin assembly and the alveolar extracellular matrix. Finally, we provide an integrated view of the dynamics of the later stages of lung development, augmenting our understanding of the cell-specific trajectories during this vulnerable developmental period.

## Results

To assess the cellular landscape during later lung development in the mouse (between E12 and P14), single-cell suspensions were generated from dissociated whole lung tissue (at least 4 mice per timepoint were pooled, and scRNA-seq was performed using the 10X Genomics Chromium platform. To specifically target the transcriptomic changes during later lung development, time points were hyper-sampled between E18 and P7 (Figure 1A). The focus of this data set is on the structural cells of the lung parenchyma; CD45+ immune cells and Ter119+ red blood cells were excluded by fluorescence-activated cell sorting (FACS) prior to sequencing library generation (Figure 1B).

**Figure 1.**
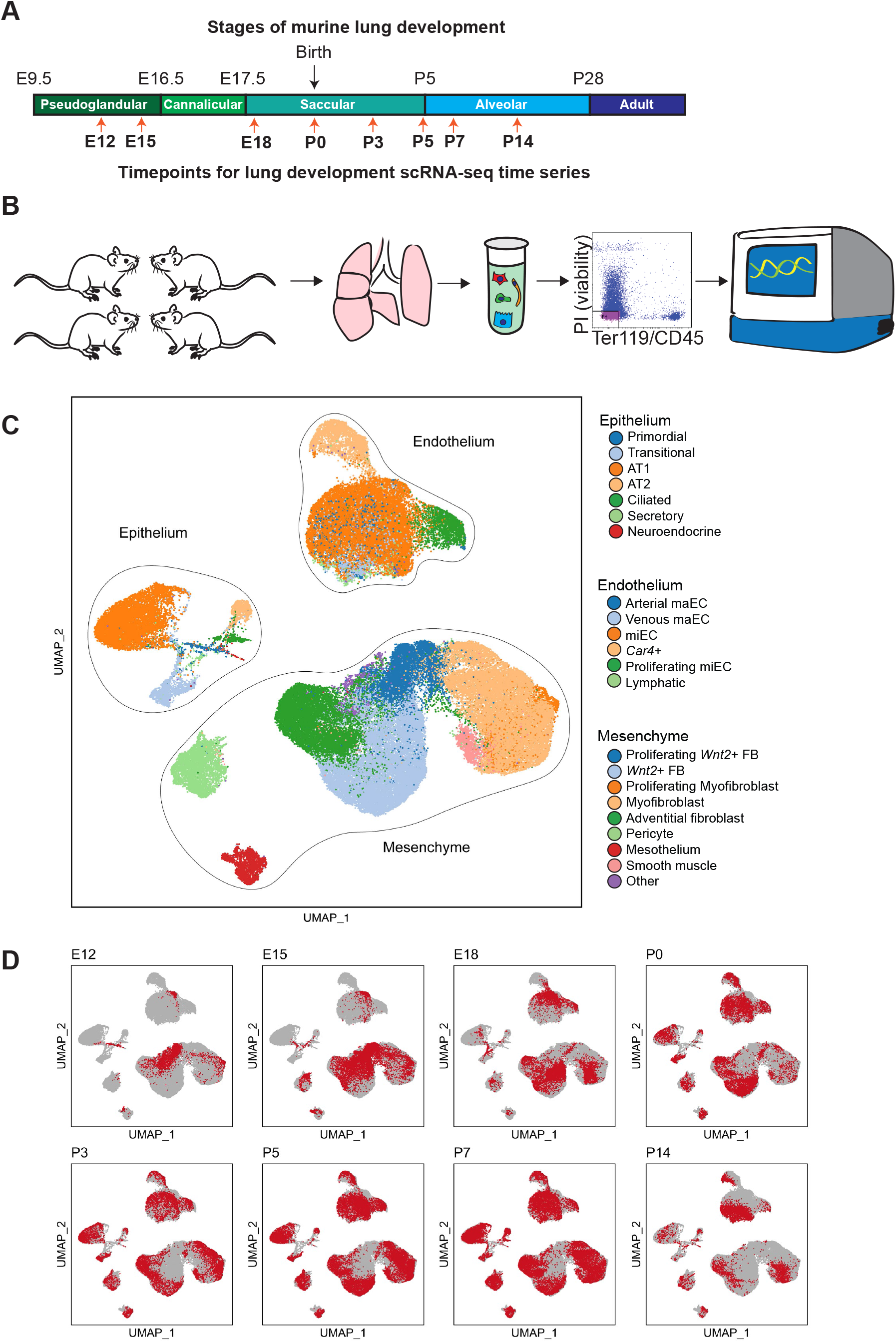
Experimental setup: capturing the process of alveolarization at a single cell resolution. **A)** Schematic of timepoints sampled for single cell RNA sequencing time series, overlayed onto histologically identified stages of murine lung development, **B)** Experimental workflow: 1. At least 4 mice were pooled per timepoint. 2. Lungs were harvested and single cell suspensions were generated by enzymatic digest. Viable, Cd45-, Ter119-cells were sorted by FACS for scRNA-seq library preparation using the 10X Genomics Chromium 5’ platform. **C)** UMAP embedding of 92,238 cells annotated by cell-type. **D)** Cells from each timepoint are highlighted in red on individual UMAP projections. Only cells from each timepoint have color, while the other cells are gray.

Jointly analyzing 92,328 single-cell transcriptomes across all time points from E12 to P14 identified 7 epithelial clusters, 6 endothelial clusters, and 9 mesenchymal clusters (Figure 1C-D, Figure S1A), with a range of 3,891 to 21,233 cells per time point. All data were jointly normalized and scaled using SCTransform function (Hafemeister and Satija, 2019) including the sequencing run as a batch variable, then jointly Uniform Manifold Approximation and Projection (UMAP)(McInnes L, 2018) embedded (Figure 1D, S1B). The epithelial, endothelial, and mesenchymal cells were then independently extracted and separately analyzed to characterize cell-type specific developmental changes over time.

### Cellular populations in the lung have temporally specified activity

Analysis of the population structure of epithelial cells (n = 11,456) from E12 to P14 identified 7 distinct clusters (Figure 1C) with notable changes over time in the relative numbers of cells in the epithelial sub-populations (Figure 2A, S2A-C). At E12, the predominant epithelial cell population is a primordial cluster identified by expression of *Mdk* (Table S1), and the relative number of cells in this population decreases significantly by the time of birth at P0. By E18, cell clusters appear that express markers of the mature lung alveolar epithelium, including a cluster of alveolar type I epithelial (AT1) cells marked by *Hopx* expression, a cluster of alveolar type II epithelial (AT2) cells marked by *Sftpc* expression, secretory cells marked by *Scgb1a1* expression, ciliated cells marked by *Foxj1* expression, and a small number of neuroendocrine cells marked by *Ascl1* expression (Table S1). Analysis of RNA velocity using scVelo(Bergen et al., 2020) was used to determine the latent time of every cell. Latent time is a proxy for cellular maturity, or an inference for how long a particular cell has been in the tissue, independent of developmental time point. Using this measure, cells that are proliferating or self-renewing exhibit decreased estimated latent time, while cells that result from a longer developmental path or existed longer within a tissue exhibit increased estimated latent time. Evaluating the expression of *Hopx*, an AT1 marker gene, we find that as latent time increases, *Hopx* increases (Figure 2B). This pattern is reversed for AT2 cells, marked by *Sftpc* expression (Figure 2C), with the highest *Sftpc* expressing cells having a lower estimated latent time, suggesting rapid AT2 biogenesis during development as well as a potential for self-renewal.

**Figure 2.**
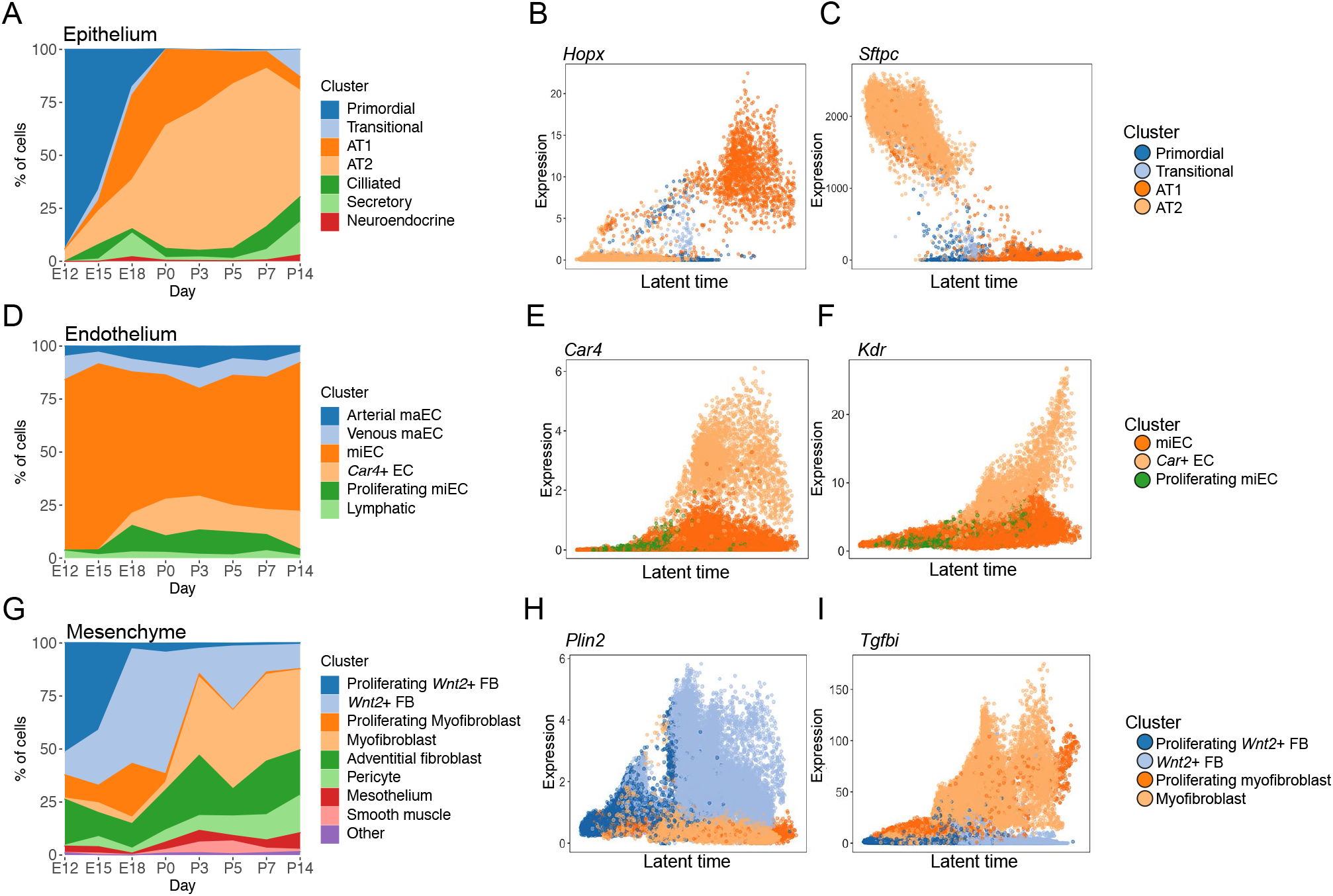
Cell populations in the lung change markedly over time. **A)** Relative proportion of epithelial cell-types as a function of time, as detected by scRNAseq. **B)** Expression of *Hopx* (AT1 marker) and **C)** *Sfptc* (AT2 marker) is plotted for each primordial, transitional, AT1, and AT2 cell. Latent time (an estimate of a cellular maturity based on RNA velocity) is plotted on the x axis with a 0 to 1 scale, and expression (as determined by scVelo) is plotted on the y axis. Colors represent cell types. **D)** Relative proportion of endothelial cell-types as a function of time, as detected by scRNAseq. **E)** Expression of *Car4* (*Car4*+ EC marker) and **F)** *Kdr* (Vegf receptor) is plotted over latent time for each microvascular endothelial cell (miEC), *Car4+* EC and proliferating miEC. **G)** Relative proportion of mesenchymal cell-types as a function of time, as detected by scRNAseq. **H)** Expression of *Plin2* (lipofibroblast marker) and **I)** *Tgfbi* (myofibroblast marker) is plotted over latent time for each proliferating *Wnt2+* FB, *Wnt2+* FB, proliferating myofibroblast, and myofibroblast.

Compared to epithelial cells, endothelial cells (n = 22,237) exhibit a more stable population structure earlier in lung development. As early as E12, there are 5 distinct endothelial clusters: 1) macrovascular arterial endothelium (arterial maEC, expressing *Vwf, Cxcl12*, and *Pcks5*), 2) macrovascular venous endothelium (venous maEC, expressing *Vwf, Vegfc*, and *Prrs23*), 3) microvascular endothelium (miEC, expressing *Gpihpb1* and *Kit*), 4) proliferating miEC (expressing *Mki67* and *Gpihpb1*), and 5) lymphatic endothelium (expressing *Flt4* and *Ccl21a*) (Figure 1C, 2D, S3A-D). At E18, a population of endothelial cells emerges that is distinguished by the expression of *Car4* (Vila Ellis et al., 2020). Analyzing expression patterns by latent time indicates that as the *Car4+* ECs have increased estimated latent time (Figure 2E) relative to other endothelial cells. Furthermore, these *Car4+* ECs express greater *Kdr* as they mature (Figure 2F).

A majority of the cells that were isolated and sequenced were pulmonary mesenchymal cells (n = 58,545). Cluster analysis indicated 9 distinct cell populations: 1) proliferating *Wnt2*+ fibroblasts, 2) *Wnt2*+ fibroblasts, 3) proliferating myofibroblasts, 4) myofibroblasts, 5) adventitial fibroblasts, 6) pericytes, 7) mesothelium, 8) smooth muscle, and 9) other (Figure 1C, 2G, S4A-C). The ‘other’ category was composed of several rare cell types including cells with expression of markers from cardiomyocytes (*Tnnt2*) or neurons (*Mpz*). While it is possible to further subdivide these clusters, parameters were chosen for clustering cell-types that were largely representative of the entire developmental window chosen. The proliferating clusters of myofibroblasts and *Wnt2*+ fibroblasts are predominantly observed in the prenatal time points, and they are distinguished by their expression of cell-cycle genes. All of the myofibroblasts (proliferating and not) express the marker genes *Acta2* and *Tgfbi*. Similarly, both of the *Wnt2*+ fibroblast clusters express *Wnt2 and Gyg* (Table S1).

In contrast to the timing of changes in the epithelium and endothelium, there is a marked shift in the relative subpopulations of mesenchymal cells between P0 and P3 (Figure 2G, S4B). The total population of all *Wnt2*+ fibroblasts decreases from 61.6% of the total mesenchymal population to 14.6%. During this time, the myofibroblasts increase from 7.6% to 38.5% of the total mesenchymal cell population. Cell-cycle analysis suggests that there is a subset of myofibroblasts that are replicating postnatally and may contribute to this population expansion (Figure S4D). Investigating expression patterns of *Plin2*, a lipofibroblast marker, demonstrates that *Plin2* expression increases in cells with increased estimated latent time (Figure 2H), and that these *Plin2+* cells are likely a subset of the *Wnt2+* fibroblasts. Expression of *Tgfbi*, a marker of myofibroblasts, emerges with a sharp inflection as a function of estimated latent time (Figure 2I).

### Trajectory analysis identifies a transitional state during alveolar epithelial differentiation

The patterns of epithelial population changes and RNA velocity suggest differentiation from both AT2 and primordial cells to AT1 cells (Figure 3A-B) in the developing lung. At E15, an epithelial population emerges that expresses markers of both AT2 cells (e.g., *Sftpc*) and AT1 (e.g., *Hopx*) (Figure S2D). These seemingly transitional cells are distinguished by the expression of several genes, including *Cdkn1a*, a cell-cycle gene (Figure 3C). While other epithelial cells outside this cluster express *Cdkn1a*, including ciliated cells, the combination of expression of markers for both AT1 and AT2 (*Aqp5, Hopx, Sfptc, Sftpa1*) as well as *Cdkn1a* was specific to this cluster (Figure 3C). Cell trajectory mapping based on mRNA-splicing (RNA velocity) analysis indicates that postnatally, these transitional cells likely arise from AT2 cells (26.9% median probability with an interquartile range of 21.8-42.8%), with a prenatal contribution from the primordial cells at early timepoints (64.2% median probability with an interquartile range of 44.2-69.0%) (Figure 3D). Trajectory mapping suggests that >95% of transitional cells become AT1 cells (Figure 3E). These cells emerge at E15, increase in relative number during between E18-P3 and remain a rare but present population through P14, a finding validated in lung tissue by quantitative RNA in situ hybridization (ISH) (Figure 3F-G, S2E). Plotting the *Cdkn1a* expression in the context of latent time further indicates that these cells have an estimated latent time that greater than AT2 cells and less than AT1 cells (Figure 3H). While expression of *Cdkn1a* (also known as p21^Cip1^) is a marker of the transitional cell population, this cluster does not have a high cell-cycle score, a multi-gene metric that identifies replicating cells (Figure S2F). Consistent with previous experimental findings (Barkauskas et al., 2013), trajectory mapping also found that AT2 cells arise from the primordial epithelium or other AT2 cells, and that AT2 cells can also give rise to AT1 cells (Figure S2G-H).

**Figure 3.**
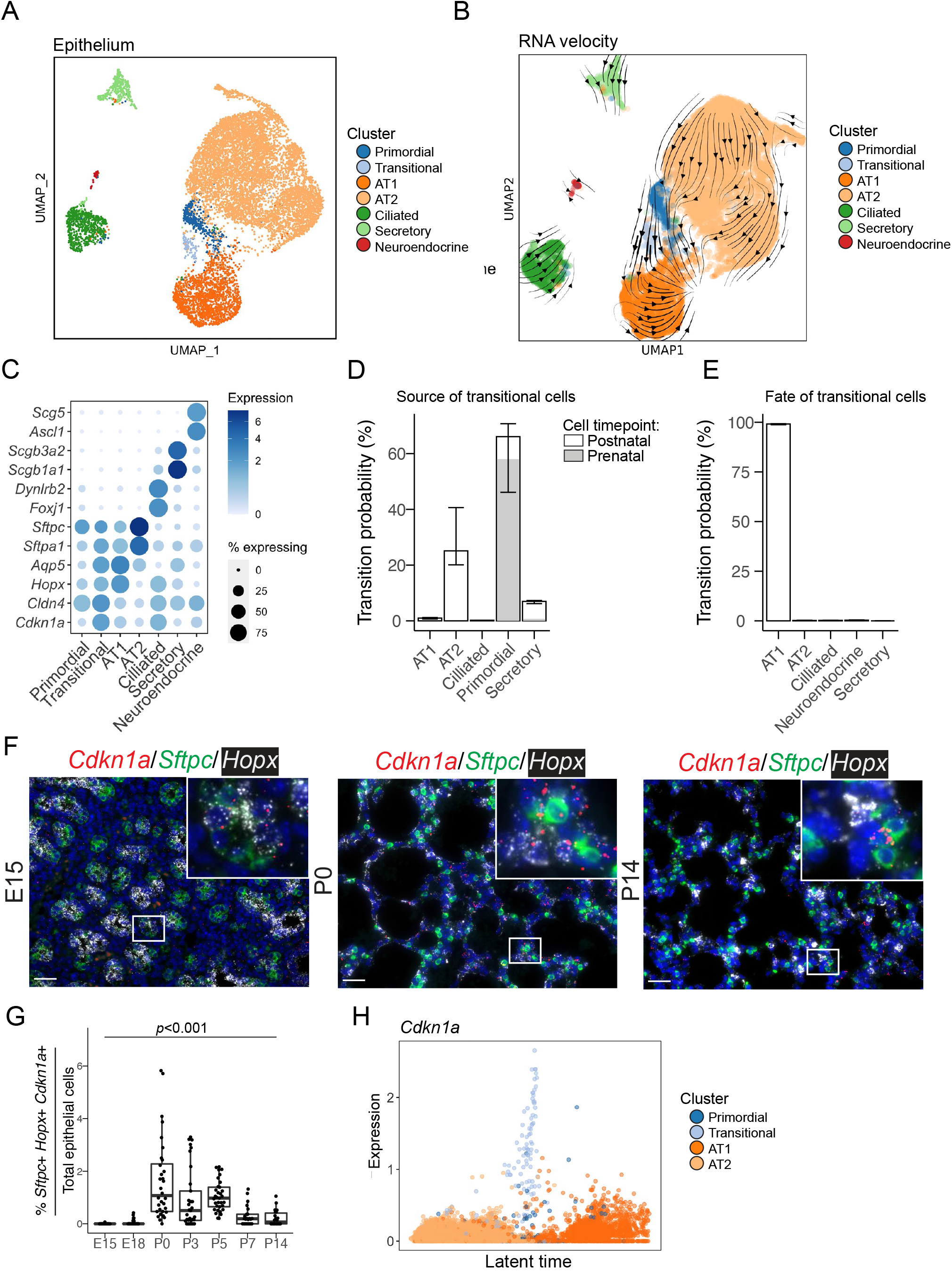
Distinct epithelial cell identities are established by E18. **A)** UMAP embedding of lung epithelial cells (n = 11,456) colored by cell type. Cells from early lungs (E12 and E15) cluster separately as indistinct epithelial cells, termed here a primordial cluster. **B)** RNA velocity vectors were calculated with scVelo and overlayed on the UMAP embedding. The thickness of the lines indicates the magnitude of the velocity. **C)** Heat map showing transitional cluster markers (*Cldn4, Cdkn1a*), AT1 markers (*Aqp5, HopX*), AT2 markers (*Sftpc, Aftpa1*), secretory cell markers (*Scgb3a2, Scgb1a1*), ciliated cell makers (*Dynlrb2, Foxj1*), and neuroendocrine cell makers (*Ascl1, Scg5*). The color intensity of the circles indicates the expression level and the size indicates the proportion of cells within a cluster that express a particular gene. **D)** Cell trajectory inference was calculated with CellRank, and the probabilities of becoming a ‘transitional’ cell are plotted. **E**) The terminal state probabilities of the transitional cells are plotted. The bars are colored based on the fraction of cells in a given cluster is present prenatally (gray) or postnatally (white). Bars represent the median and error bars represent the interquartile range. **F)** RNA in situ hybridization (ISH) of *Cdkn1a* (red, transitional cell marker), *Sftpc* (green, type II cell marker), and *Hopx* (white, type I cell marker) at selected timepoints. Scale bar = 25 µm. A complete series is presented in Figure S2E. **G)** Quantification of the ISH using HALO for percent of transitional cells (*Sftpc+Hopx+Cdkn1a+*) cells over total epithelial cells. A non-parametric Kruskal-Wallis test was performed (*p* < 0.001). **H)** Expression of *Cdkn1a* (transitional cell marker) is plotted for each primordial, transitional, AT1, and AT2 cell. Latent time is plotted on the x axis with a zero to 1 scale, and expression (as determined by scVelo) is plotted on the y axis.

### Early specification is characteristic of the developing lung endothelium

While the lung endothelium has a relatively stable population structure from E12 to P14, *Car4+* endothelial cells emerge at E18 (Figure 2D, 4A). RNA velocity analysis and trajectory inference of the endothelial cells indicated that the source of the *Car4*+ cells is likely the populations of miEC cells (67.8% median probability with an interquartile range of 66.3-69.0%) and proliferating miEC cells (26.1% with an interquartile range of 24.9-26.3%) (Figure 4B-C), consistent with observations in the adult mouse lung (Gillich et al., 2020). As anticipated, velocity analysis of the endothelium suggests that miECs are capable of self-renewal (Figure S3F), confirming previous work (Kawasaki et al., 2015). This is also congruent with the finding that there is a cluster of miECs with a high cell-cycle score, indicating proliferating miECs (Figure S3G). Furthermore, using trajectory mapping, we now define a small subset of miEC cells with an elevated *Car4*+ fate probability. This population can also be identified as miEC cells that have increased latent time compared to nearby cells in the UMAP (Figure 4D). These cells begin to express genes that are markers of the *Car4*+ population, including *Igfbp7, Kitl*, and *Tbx2* in addition to *Car4* expression (Supplemental Table 2). Furthermore, as the *Car4*+ cells mature along the predicted trajectory defined by latent time, the expression of *Kdr*, a VEGF receptor, increases (Figure 2F, 4E). Spatial examination of *Car4* expression in lung tissue confirms that beginning at E18, there is relatively consistent population of *Car4*+ endothelial cells found in the alveoli by quantitative RNA ISH (Figure 4F-G, S3E).

**Figure 4.**
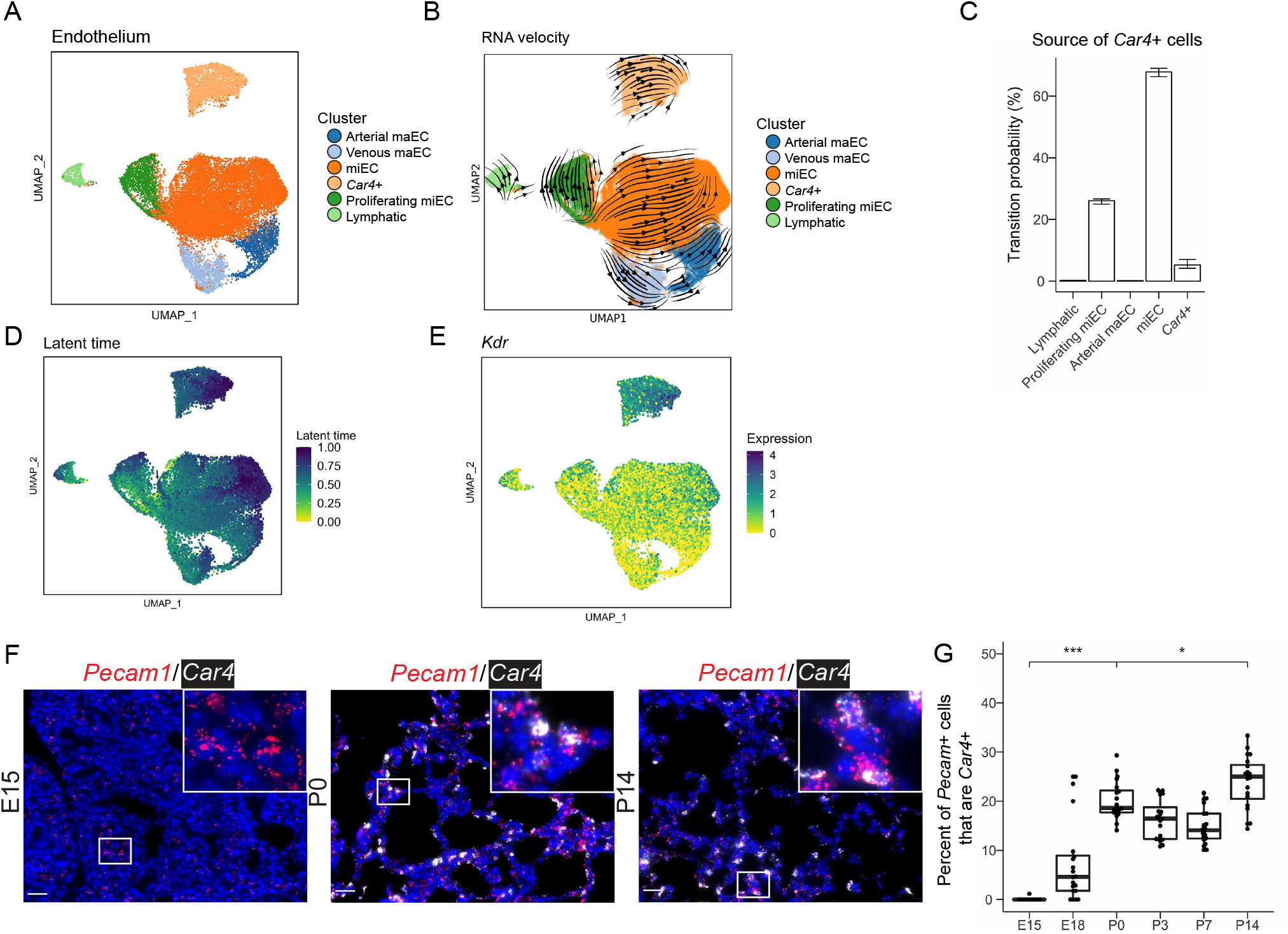
Endothelial cells have specification by E15, with an emergence of *Car4*+ cells by E18. **A)** UMAP embedding of lung endothelial cells (n = 22,237) colored by cell type. **B)** RNA vectors of the endothelial cells were calculated with scVelo and overlayed on the UMAP embedding. **C)** Cell fate prediction was calculated with CellRank, and the predicted source of the *Car4+* cells was plotted. Bars represent the median and error bars represent the interquartile range. **D)** UMAP embedding of endothelial cells colored by cell latent time where darker colors indicate greater latent time. The arrow indicates the small group of miEC cells that have high latent time and are likely to transition to *Car4*+ cells. **E)** UMAP embedding of endothelial cells colored by *Kdr* expression where darker colors indicate greater expression. **F)** RNA in situ hybridization (ISH) of *Pecam1* (red, endothelial cell marker), and *Car4* (white, *Car4+* cell marker). Scale bar = 25 µm. A complete series is presented in Figure S3E. **G)** Quantification of the ISH was performed using HALO and the fraction of *Pecam*+ cells that are *Car4*+ was plotted. Boxplots represent the summary data and each dot represents the percentage of *Pecam*+*Car4*+ / *Pecam*+ cells per image (n = 20 images per time point). Increases were observed from E15 to P0 and from P0 to P14. (* *p* < 0.05, *** *p* < 0.001 by Mann–Whitney U test).

### Mesenchymal cells retain distinct identities but alter expression patterns and abundance during development

Compared to epithelial and endothelial cells, mesenchymal cells undergo more substantial population shifts after birth, suggesting postnatal cellular replication and transition (Figure 5A-B, S4A-C). Quantitative ISH was used to stain and identify the myofibroblasts with marker *Tgfbi*+ and *Wnt2*+ fibroblasts in the developing lung (Figure 5C, S4E). Examination of lung tissue with these markers found that most of the cells in the E15 lung are *Wnt2*+ fibroblasts (Figure 5C, S4E) with a dramatic redistribution of the myofibroblasts (marked by *Tgfbi* expression) from large clusters of *Tgfbi*+ cells at E15 to a spread out population of myofibroblasts at E18 and progressing postnatally to P0. Spatial analysis of at least 3,200 cells per time point using the Clark and Evans aggregation index supported the observation that the greatest organizational changes occurred between E15 and P0, with the *Wnt*2+ fibroblasts becoming more randomly distributed around alveoli and the myofibroblasts becoming less clustered with time (Figure 5D). Analysis of RNA velocity and cell trajectory mapping suggested that the prenatal proliferating myofibroblasts predominately self-renew, but also have a possibility of becoming *Wnt2*+ fibroblasts. In contrast, the proliferating *Wnt2*+ fibroblasts almost exclusively become mature *Wnt2*+ fibroblasts (Figure 5E-F). Cell fate analysis did not identify a common mesenchymal progenitor population, suggesting that some mesenchymal fate specification occurs earlier than E12.

**Figure 5.**
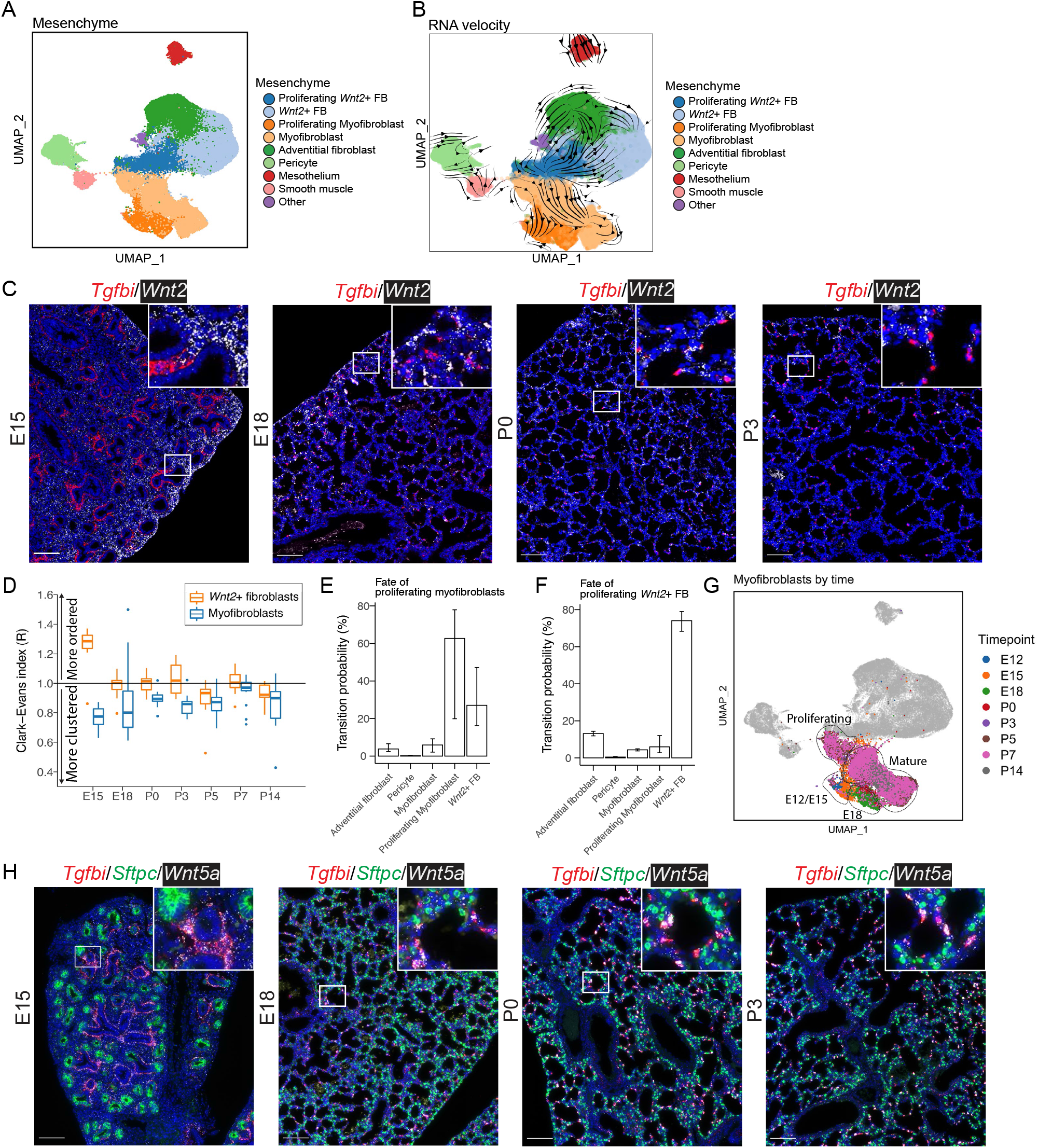
The mesenchymal cell population undergoes dynamic population shifts between P0 and P3. **A)** UMAP embedding of lung mesenchymal cells (n = 58,545) colored by cell type. **B)** RNA velocity vectors overlayed on the UMAP depict inferred differentiation/maturation trajectories. **C)** RNA in situ hybridization (ISH) of *Tfgbi* (red, myofibroblast marker) and *Wnt2* (white, *Wnt2+* fibroblast marker). Images are stitched scans, scale bar = 100 µm. A complete time series of representative images is presented in Figure S4E. **D)** HALO was used to analyze the results of the *Tgfbi* and *Wnt2* ISH and assign cell identities based on marker expression. To determine the degree of clustering, a Clark-Evans index was calculated from individual cells and aggregated by image. Values greater than 1.0 indicate that the cells are more ordered (regularly spaced), while values less than 1.0 indicate that the cells are more clustered. **E)** Cell fate prediction was calculated with CellRank, and the predicted fate of the proliferating myofibroblasts and **F)** *Wnt2*+ fibroblasts was plotted. Bars represent the median and error bars represent the interquartile range. **G)** A UMAP embedding of myofibroblasts colored by the age of the mouse, all other cell types are gray. Sub-clusters of the myofibroblasts are roughly indicated in an outline. **H)** RNA ISH of *Tgfbi* (red, myofibroblast marker), *Sftpc* (green, type II cell marker), and *Wnt5a* (white). Images are stitched scans, scale bar = 100 µm. A complete time series of representative images is presented in Figure S5D.

More detailed analysis of the expression patterns in myofibroblasts shows that there is clustering by time point, where the E12 and E15 myofibroblasts cluster together, separately from the E18 myofibroblasts, which have increased expression of *Wnt5a, Pdgfra* and *Enpp2* (Figure 5G, Figure S5A, Table S3). Postnatally, there are two distinct groups of myofibroblasts, one group is marked by a high cell-cycle score (Figure S4D) and is likely a persistent group of proliferating myofibroblasts, while the other group appears to be the mature stable myofibroblasts. It was also noted that while *Wnt2*+ fibroblasts all broadly expressed *Wnt2*, the myofibroblasts (starting at E18) expressed *Wnt5a* (Figure S5A-C, Table S3). Quantitative ISH supports this finding, as >90% (median) of *Tgfbi*+ myofibroblasts co-express *Wnt5a* (Figure 5H, Figure S5D-E).

### During the later stages of lung development, alveolar Type 1 cells express genes involved in elastin assembly and extracellular matrix organization

As an application of this cellular atlas, we explored the expression profile of AT1 cells during later lung development. Previous work has suggested that proper elastin assembly and generation of the basement membrane and extracellular matrix (ECM) are critical steps during the saccular and alveolar stages of lung development (Bourbon et al., 2005). We observed that AT1 cells highly express *Fbln5* (fibulin-5) (Figure 6A), which contributes to elastin assembly by binding tropoelastin and fibrillin-1 and by cross-linking enzymes required in this process. AT1 cells also expressed components of the basement membrane, including multiple components of type IV collagen and laminins. Laminins are trimeric ECM proteins, comprised of an alpha, beta, and gamma subunit, found in the lung basement membrane, but the granular timing of laminin gene expression and the cellular source is poorly defined. Our analysis found expression of laminin-332 (made of α3, β3, and γ2), a laminin isoform present in the adult alveolar basement membrane), is high at E18 in AT1 cells and continues through development. Spatially, AT1 cells had a peak of *Fbln5* expression between P0-P5 (Figure 6A), and RNA ISH confirmed the colocalization of AT1 marker gene *Hopx* expression with *Fbln5* across our time series. Quantification of RNA ISH by automated image analysis (Halo, Indica labs) demonstrated increased expression of *Flbn5* by *Hopx+* cells, with a peak between P3-7 (Figure 6B-D, S6A). Consistent with this finding, *Fbln5* expression was found to increase in AT1 cells with greater estimated latent time (Figure 6D). The expression profile of elastin assembly components and ECM in AT1 cells is unique among epithelial cells during development, with AT2 cells having little detectable expression of these genes (Figure S6B).

**Figure 6.**
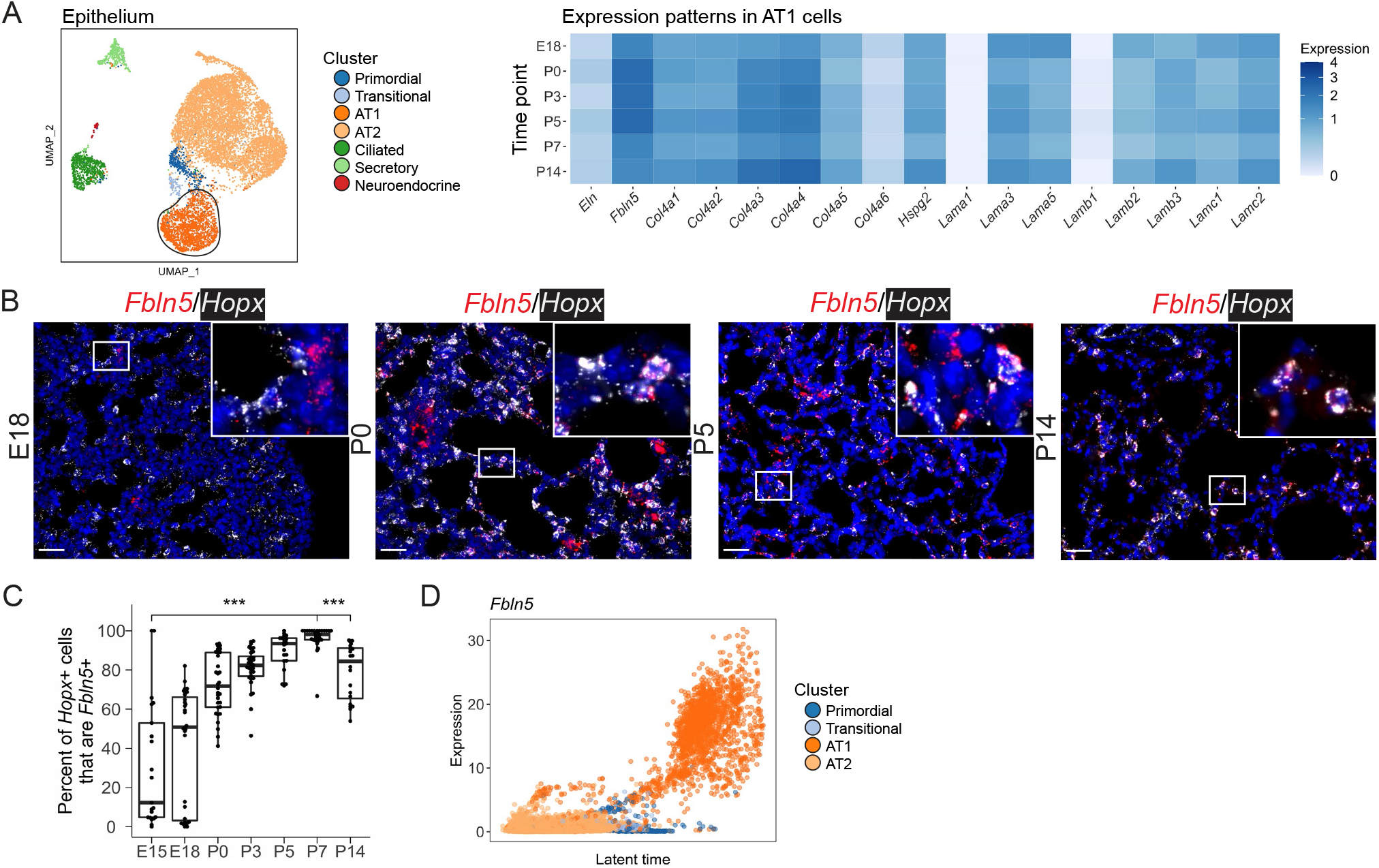
Epithelial alveolar type I (AT1) cells express components of elastin and the basement membrane. **A)** Expression patterns in AT1 cells were analyzed over time. Heatmaps represent average expression in a given cluster, where darker colors indicate more expression. B) RNA in situ hybridization (ISH) of *Fbln5* (red), and *Hopx* (white, epithelial type I marker). Scale bar = 25 µm. A complete series is presented in Figure S6A. **C)** Quantification of the ISH was done using HALO. Boxplots represent the summary data, and each dot represents the percentage of *Fbln5+Hopx*+ / *Hopx*+ cells per image (n > 20 images per time point, *** *p* < 0.001 by Mann–Whitney U test). **D)** Plot of *Fbln5* expression (as determined by scVelo) by latent time in primordial, transitional, AT1, and AT2 cells, where the x-axis is on a scale from 0 to 1.

## Discussion

This study creates a cellular atlas of epithelial, mesenchymal, and endothelial cells of the later stages of lung development, identifies new temporal relationships between these cell types, and defines the relative populations of major subtypes of these cells during the later stages of lung development (Figure 7). We have applied new analysis paradigms to understand the trajectory and fate prediction of these cell types, contextualizing previous findings and discovering new features of developmental lung biology. These data now provide a developmental roadmap that identifies the stages of cell-type commitment in the lung, and a resource to define the cellular sources of important developmental factors.

**Figure 7.**
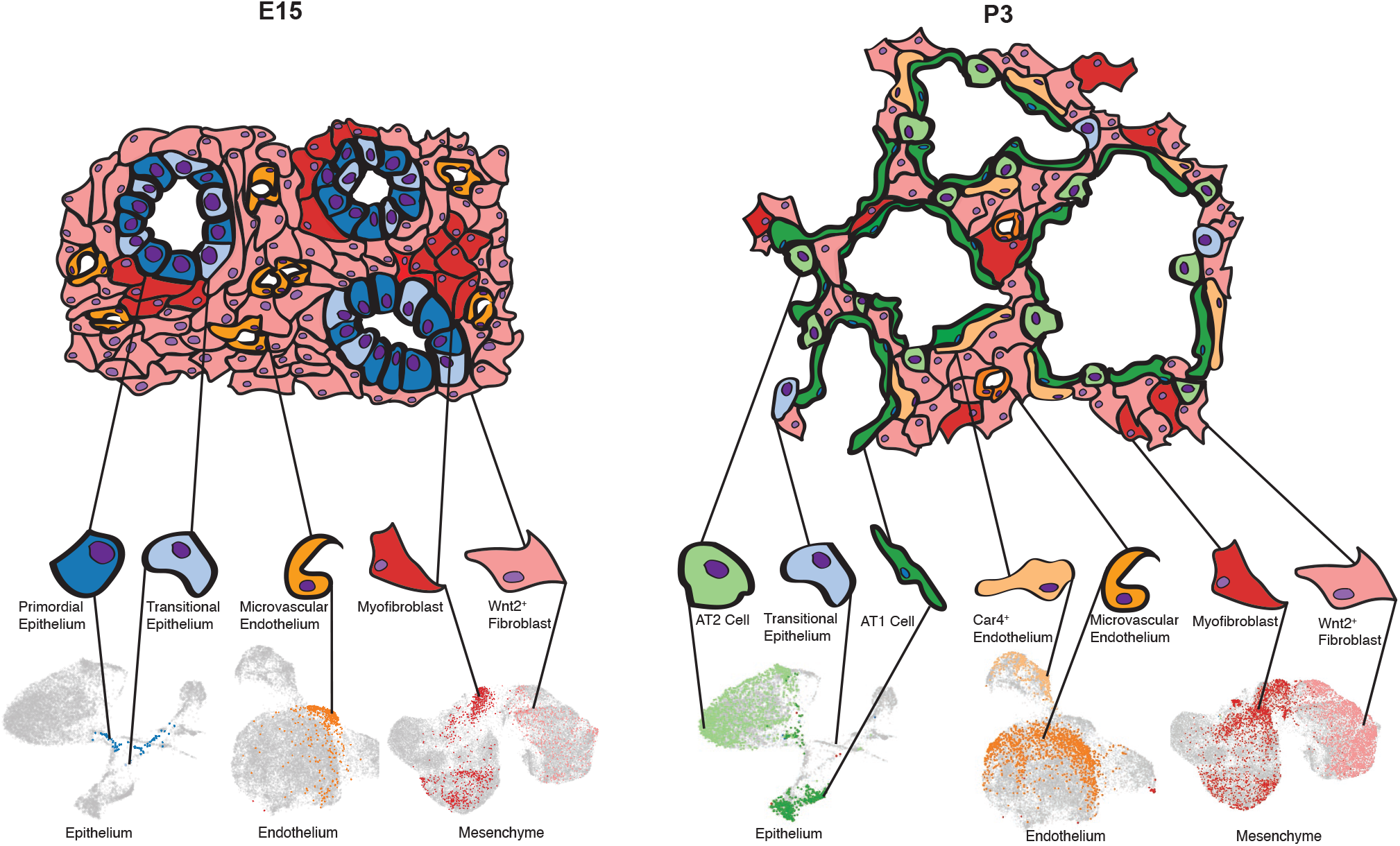
Functional development of the lung is a result of spatial and temporal organization of multiple cell types. A model of the developing lung at two representative timepoints is presented. At E18, the lung epithelium is composed of indistinct cell types that are very rare or absent in adulthood. The prenatal endothelium in the lung parenchyma is composed of microvascular endothelial cells, while the majority of the cells in the remaining space of the lung are *Wnt2*+ fibroblasts, with interspersed clusters of myofibroblasts. By P3, the cell types observed in mature lungs are present, including AT2, AT2, and transitional epithelial cells. The microvascular endothelium has differentiated into *Car4*+ cells, alongside the microvascular endothelium, and the mesenchymal cells have undergone extensive spatial reorganization.

While the lung epithelium at E12 and E15 is primarily composed of indistinct “primordial” cells, a few type I and type II cells can be observed as early as E15, suggesting a relatively early fate commitment for at least a subset of alveolar epithelial cells, consistent with some previous findings (Frank et al., 2019). Ongoing AT1 differentiation from AT2 cells occurs during alveologenesis, suggesting multiple differentiation trajectories contribute to the maturing AT1 population (Desai et al., 2014). The transitional epithelial cells, which we identify as *Cdkn1a+*, are similar to epithelial cells previously described in lung organoids as the pre-alveolar type-1 transitional cell state (PATS) (Kobayashi et al., 2020) and the damage associated transient progenitors (DATP) (Choi et al., 2020), which arise from adult AT2 cells in the setting of injury. Similar to the DATPs after injury, our trajectory analysis now establishes that these *Cdkn1a*+ transitional cells arise from AT2 cells postnatally, and that >95% of these neonatal transitional cells ultimately become AT1 cells. Interestingly, in this dataset, it appears that prenatally, these transitional cells primarily arise from early primordial epithelial cells, and that a small number of primordial epithelial cells (0.2% of total epithelial cells) persist as late as P7. In the adult mouse, DATP appear to be essential for regeneration of the lung after injury (Choi et al., 2020). Future work is needed to elucidate the primordial and transitional cell populations in the setting of neonatal injury and their governing mechanisms. While Wnt responsiveness has been shown to be relevant to regeneration and repair in the adult lung (Zacharias et al., 2018), in this developmental dataset, both primordial and transitional cell populations lack detectable expression of *Axin2*. This suggests that Wnt signaling is not a significant feature of cells in these states during normal development, though Wnt may still play a role up or downstream of these cell states, or in the setting of injury.

This time-series demonstrates that the recently identified *Car4*+ endothelial cells (Gillich et al., 2020; Niethamer et al., 2020; Vila Ellis et al., 2020) emerge at E18, and these *Car4*+ cells persist in relatively stable numbers though P14. Trajectory analysis identified that *Car4*+ cells arise from the miEC population, and the probability of cells differentiating into *Car4*+ endothelial cells appears to be correlated with the expression of *Igfbp7, Kitl*, and *Tbx2*. These specialized endothelial cells are closely related to AT1 epithelial cells spatially and temporally. We observed that the *Car4*+ cells appear at E18, at the same time as the neighboring AT1 cells emerge. With time, *Car4*+ endothelial cells begin to express higher levels of VEGF-receptor *Kdr4*, as *Vegfa* levels concurrently increase in AT1 cells. This is consistent with the previous finding that Vegf is an essential factor for the formation of *Car4*+ endothelial cells that are found in association with alveolar structures (Vila Ellis et al., 2020).

The greatest diversity of cell populations and dynamics during the later stages of lung development can be found in the pulmonary mesenchyme. Prenatally, the predominant population expresses *Wnt2*. Later in development this population also expresses other markers associated with lipofibroblasts (e.g., *Plin2, Tcf21, Gyg*) (Endale et al., 2017). Just prior to birth, the myofibroblast population begins to expand and express a different Wnt ligand, *Wnt5a*. The spatial and temporal shifts of these two fibroblast populations are highly dynamic between E15 and P5. Spatial analysis of RNA ISH shows that at E15, both populations are highly clustered. During sacculation and early alveologenesis, both groups of fibroblasts spread out and intercalate next to each other in the crests of alveoli, with *Wnt5a*+/*Tgfbi*+ myofibroblasts present in a very small number of cells in each alveolus. Our prior work has demonstrated a pathologic role for excessive *Wnt5a* expression by mesenchymal cells during neonatal hyperoxia injury (Sucre et al., 2020), and we speculate that increased *Wnt5a* expression by the myofibroblast population or expansion of this population after injury is a key driver of abnormal development observed after neonatal hyperoxia exposure.

The *Wnt2*+ populations we observed pre- and postnatally differ significantly from previously described *Wnt2*+ fibroblasts in adult lungs (Zepp et al., 2017). In the adult population, it has been reported that >90% of *Wnt2*-lineage-labeled fibroblasts express *Pdgfra*. In our developmental dataset, the opposite appears to be true, with *Pdgfra* expressed in higher levels in the *Tgfbi*+ myofibroblast population. However, our trajectory analysis found that some of the proliferating myofibroblasts have the predicted potential to become *Wnt2+* fibroblasts. This suggests that while there may be early overlap in these populations, the gene expression profiles after commitment to a specific cell type are distinct. In addition to the age of the mice studied, there are technical reasons why our observations may differ from prior work. Our scRNA-sequencing looked at all CD45-, viable cells in the lung, without the use of fluorescent reporters to enrich for a specific target population. While this allowed for a global, unbiased look at the structural cells in the lung, the ability to detect very rare populations of cells within the lung is decreased. We observe expression of previously described *Lgr6*+ cells in the myofibroblast population beginning at E12, but rarer populations, such as the *Lgr5*+ mesenchymal cells (which comprise 1-2% of pulmonary mesenchymal cells in the adult lung) (Lee et al., 2017) are not present in sufficient quantity to be detected by our analysis. In addition to the tissue validation we have done with RNA ISH, future work with reporter lines could certainly be used for lineage tracing and confirmation of our findings, though these methods are not without their limitations related to fidelity and may overestimate or underestimate the number of cells in a particular population (Hogan, 2018).

Profiling the developing lung over time has allowed greater insight into specific cellular transitions than studying a single timepoint. Specifically, by examining a dataset that is composed of immature cells, mature cells, and cells in transition, it is possible to make inferences about the trajectory of individual cells using scRNAseq. However, the approach of scRNAseq does not facilitate tracking live cells over time. There are other notable limitations to this study, most obviously that it excludes all CD45+ immune cells. The resident and circulating immune cells of the lung have been shown to contribute significantly to both lung development and injury response (Stouch et al., 2016). In order to increase the numbers of epithelial, mesenchymal, and endothelial cells that were sequenced and analyzed, we selected out this important population. Other limitations of this study include the biases inherent in scRNAseq, including that the single cell dissociation may bias toward recovery and analysis of certain populations. While the actual proportions of cell subpopulations in the lung may be different, we anticipate that the relative changes in population over time are preserved. In addition, our trajectory inferences about RNA velocity may be reversed if transcript abundance is regulated by RNA degradation; the correct prediction of established lineage relationships in the epithelium and endothelium provides confidence that this concern is less likely a major confounder. Despite these limitations, this study has several strengths, including the large number of cells sequenced (92,238) across developmental timepoints, the inclusion of at least 4 mice per timepoint, and application of advanced analysis techniques to allow for the study of trajectory and fate mapping across developmental time.

We have applied the findings of this developmental cell atlas to study the extracellular matrix gene expression profile of AT1 cells. Previous work has demonstrated that proper elastin assembly is critical for alveologenesis (Benjamin et al., 2016). We identified that AT1 cells express high levels of *Flbn5*, which is essential for elastin crosslinking and proper assembly. Laminin is one of the principal components of the lung basement membrane and is required for lung branching morphogenesis (Nguyen et al., 2005; Schuger et al., 1990). Others have previously reported different laminin isoforms expressed by the fetal lung than in the adult alveolus; with laminin-111 and −511 expressed early and laminin-332 and −511 in the adult alveolar basement membrane (Miner et al., 1995; Miner et al., 1997; Nguyen and Senior, 2006). Our data precisely define the appearance of laminin-332 at E18 and identify AT1 cells as a cellular source of this laminin beginning at E18. While AT1 cells have been previously characterized as providing a functional surface for gas exchange and for regulating permeability and tight-junctions of the alveolus (Warburton et al., 2010), taken together, these findings suggest AT1 cells are a cellular source of critical ECM components as the developing lung transitions from the saccular and alveolar stages. Future work with *in vivo* models is needed to characterize this newly identified function of AT1 cells.

The developmental stages of the lung have been previously defined descriptively by histologic features (Warburton et al., 2010). This categorization often results in regarding the function and identity of cell types within the border zone of each stage. In contrast to a histologic strategy of staging, we approached development from the perspective of cellular function as defined by the transcriptome. In aggregate, we found that different cell types commit to change asynchronously during development, suggesting that the timing of the saccular-alveolar transition is fluid and cell-type specific. Combined, the dynamic expansions and contractions of cell populations, including epithelial specification around E15, the appearance of *Car4*+ endothelial cells around E18, and an expansion of myofibroblasts around P3, indicate that there is tremendous coordination and interaction between epithelial, mesenchymal, and endothelial cells at every time point. This granular single cell atlas of the developing lung provides insight into the relevant populations present between E12 and P14 and provides a foundation for future mechanistic studies into pathways governing the coordinated cell-cell communication across this specialized developmental stage.

## Acknowledgements

This work was supported by NIH K08143051 (JMSS), K08HL130595 (JAK), R01HL145372 (JAK/NEB), P01HL092470 (TSB), K08HL127102 (EJP), R03HL154287 (EJP), K08HL133484 (JTB), and The Francis Family Foundation (JAK and JMSS). Flow Cytometry experiments were performed in the VMC Flow Cytometry Shared Resource which is supported by the Vanderbilt-Ingram Cancer Center (P30 CA68485) and the Vanderbilt Digestive Disease Research Center (DK058404).

## Author Contributions

Conceptualization: NMN, EJP, JTB, JAK, JMSS, Data curation: NMN, JAK, JMSS, Formal analysis: NMN, BAS, JAK, ACH, NEB, JMSS, Funding acquisition: JAK, JMSS, Investigation: NMN, EJP, JTB, BAS, ACH, CSJ, CJT, PG, DN, BKM, JMSS, JAK, Methodology: EJP, BKM, JAK, BAS, NMN, JMSS, NEB, Project Administration: CSJ, PG, Software: NMN, BAS, ACH, JAK, Validation: NMN, EJP, JTB, JAK, JMSS, Visualization: NMN, JAK, ACH, BAS, JMSS, Writing (original draft): NMN, EJP, JTB, JAK, JMSS, Writing (review and editing): NMN, EJP, JTB, BAS, SHG, TSB, NEB, JAK, JMSS

## Materials and Methods

### Animal sample collection and care

C57BL/6 mice were used for all experiments in this study. Timed matings were performed as previously described (Plosa et al., 2020) and mice were sacrificed at postnatal day (P) 0, P3, P5, P7, or P14 for single cell RNA sequencing or lung block fixation. Embryonic day (E) 12, E15 and E18 lungs were isolated by removing pups from the mouse uterus and isolating lung tissue. E12, E15, E18, P0, P3, and P5 lungs were fixed in formalin. P7 and P14 mouse lungs were inflation-fixed by gravity filling with 10% buffered formalin and paraffin embedded. All animal work was approved by the Institutional Animal Care and Use Committee of Vanderbilt University (Nashville, TN) and was in compliance with the Public Health Services policy on humane care and use of laboratory animals.

### Single cell data collection

Sample collection and single cell sequencing was performed as previously described(Schuler et al., 2020). Briefly, lung lobes were harvested, minced, and incubated for 30 minutes at 37°C in dissociation media (RPMI-1640 with 0.7 mg/ml collagenase XI and 30 mg/ml type IV bovine pancreatic DNase). After incubation, tissue was disassociated into single cell suspension by passage through a wide bore pipet tip and filtration through a 40 µm filter. Single cell lung suspension was then counted, aliquoted, and blocked with CD-32 Fc block (BD cat #553142) for 20 minutes on ice. After 2% FBS staining buffer wash, cells were incubated with the conjugated primary antibodies anti-CD45 (BD cat # 559864) and anti-Ter119 (Biolegend cat# 116211). For some of the P7 libraries, epithelial enrichment was performed on some samples by incubation with Epcam antibody (BD cat # 563477). In the same manner, fluorescence minus one controls were blocked and stained with the appropriate antibody controls. Cells from individual mice were incubated with identifiable hashtags, resuspended in staining buffer, and treated with PI viability dye. CD45 negative, Ter119 negative, viable cells were collected by fluorescence associated cell sorting using a 70 µm nozzle on a 4-laser FACSAria III Cell Sorter. For epithelial enriched samples, Epcam+ cells were sorted. Both single and fluorescence-minus-one controls were used for compensation.

### scRNA-seq library preparation and next-generation sequencing

ScRNA-seq libraries were generated using the 10X Chromium platform 5’ library preparation kits (10X Genomics) following the manufacturer’s recommendations and targeting 10,000 − 20,000 cells per sample. Next generation sequencing was performed on an Illumina Novaseq 6000. Reads with read quality less than 30 were filtered out and CellRanger Count v3.1 (10X Genomics) was used to align reads onto mm10 reference genome.

### Analysis of single cell sequencing data

Ambient background RNA were cleaned from the scRNA-seq data with “SoupX” (Young and Behjati, 2020) as described previously (Schuler et al., 2020) using the following genes to estimate the non-expressing cells, calculate the contamination fraction, and adjust the gene expression counts: *Dcn, Bgn, Aspn, Ecm2, Fos, Hbb-bs, Hbb-bt, Hba-a1, Hba-a2, Lyz1, Lyz2, Mgp, Postn, Scgb1a1*. Quality filtering was then used to remove cells with > 10% mitochondrial mRNA, < 0.5% 10% mitochondrial mRNA, and cells with < 700 detected genes.

Preliminary data analysis, dimensionality reduction, clustering, visualization, and grouping of cells into broad categories (epithelial, endothelial, and mesenchymal cells) was performed using Seurat v3.2.2 and SCTransform (Hafemeister and Satija, 2019; Stuart et al., 2019), with manual inspection of the expression patterns of the marker genes: *Hbb-bs, Epcam, Foxj1, Scgb1a1, Scgb3a2, Abca3, Hopx, Col1a1, Dcn, Lum, Acta2, Wnt2, Wnt5a, Lgr6, Pdgfra, Pdgfrb, Cspg4, Wt1, Pecam1, Ccl21a, Vwf, Nrg1, Plvap, Car4, Mki67, Tnnt2*, and *Mpz*. Cell clusters with containing non-physiologic marker combinations, i.e. Epcam+/Pecam1+, were dropped at this stage. SCTransform was run with each sequencing reaction (corresponding to an individual timepoint) as a batch variable, and with the percentage of mitochondrial RNA as a regression variable.

Further data cleaning was done to remove gene counts for *Gm42418*, which is likely a rRNA (Kimmel et al., 2019). After removal of *Gm42418*, genes with expression in less than 10 cells across the dataset were removed from further analysis. SCTransform (version 0.3.1.9002) was used with glmGamPoi (version 1.2.0) (Ahlmann-Eltze and Huber, 2020) to normalize the data prior to PCA, UMAP, and cell clustering individually for each broad cell category (epithelial, endothelial, and mesenchymal). Cell clusters for each broad category were identified using the marker genes outlined in the supplemental figures in addition to marker gene detection with MAST (Finak et al., 2015). Data integration was performed on the second P7 sample using the SCTransform integration workflow, which is based on cross correlation analysis of gene expression Pearson residuals. After annotation of all cell clusters, the broad tissue categories were merged, and dimensionality reduction, clustering, visualization were conducted. All charts and heatmaps as part of the scRNAseq analysis were generated with ggplot2, and all parts of the analysis was run in R 4.0.2.

Predictions of RNA velocity were made using a velocyto, scVelo, cellrank pipeline. Spliced and unspliced mRNA counts were determined with velocyto (version 0.17.17) (La Manno et al., 2018) and analyzed using scVelo (version 0.2.2) (Bergen et al., 2020). Cellrank (vesion 1.1.0) (Lange M, 2020)was then used to infer the initial and terminal cell states. Cell cycle scores were calculated using scVelo. All packages were run in Python 3.8.5. Specific parameter settings, a complete collection of all package versions, and code for all steps of the analysis is available at https://github.com/SucreLab/LungDevelopment.

## Data availability

All sequencing data is available at the NCBI GEO database under accession numbers GSE160876 and GSE165063.

## RNA in situ hybridization

RNAScope technology (ACDBio) was used to perform all RNA *in situ* hybridization (RNA ISH) experiments according to manufacturer’s instructions. RNAScope probes to the following mouse genes were used: *Sftpc, Hopx, Cdkn1a, Wnt2, Tgfb1, Dcn, Pecam1, Car4, Flbn5*. Positive control probe (*PPIB)* and negative control probe (*DapB)* were purchased from the company and performed with each experimental replicate, with representative data shown (Figure S6C).

## Image acquisition and analysis

Fluorescent images were acquired using a Keyence BZ-X710 or BZ-X810 with BZ-X Viewer software with 40X objective. Wavelengths used for excitation included 405, 488, 561, and 647nm lines. Automated image analysis was performed with Halo software (Indica Labs); an example of cellular segmentation and determination of co-localization is demonstrated in Figure S6D.

## Supplementary Figures

**Figure S1.**
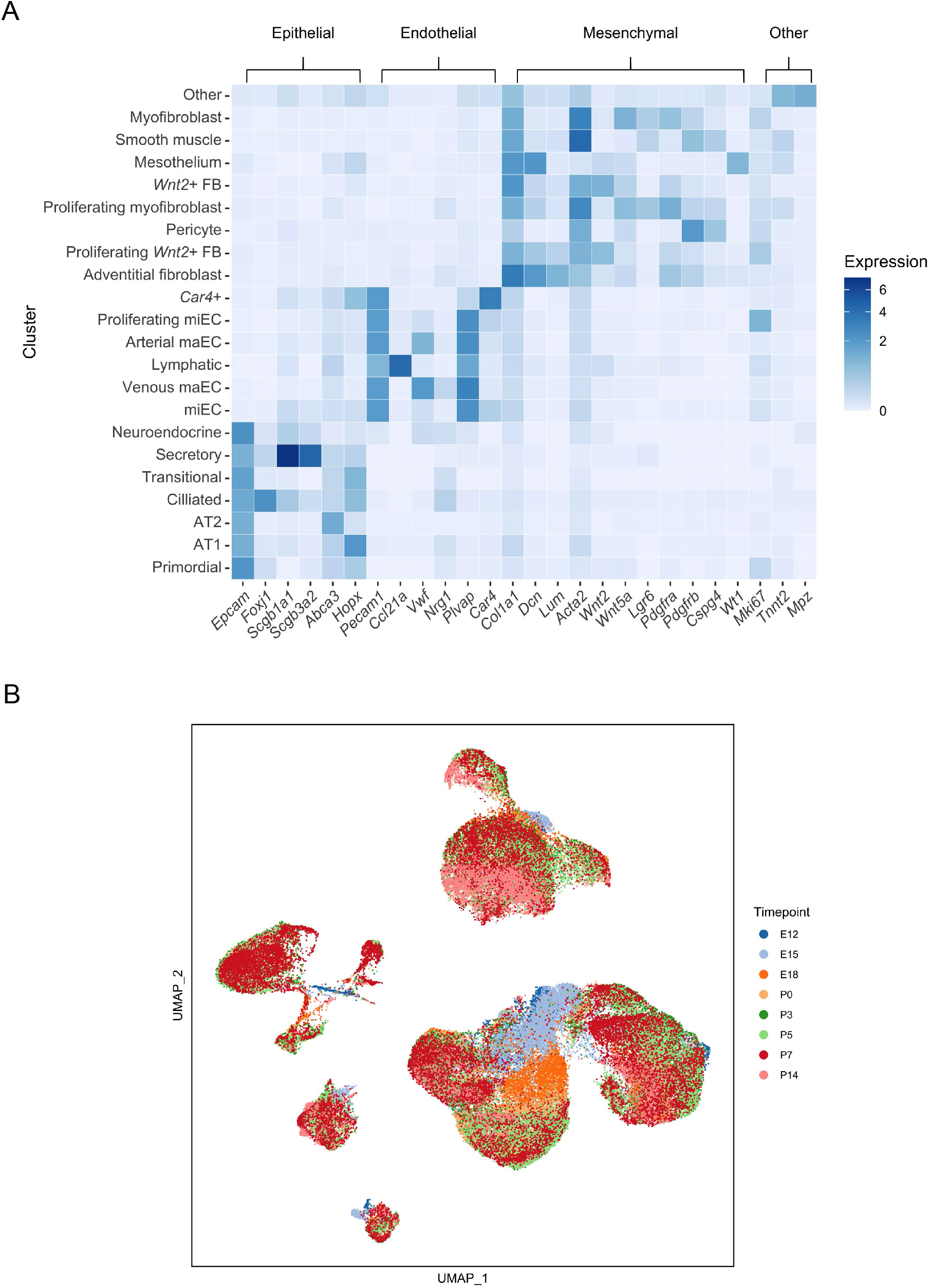
Preliminary analysis of the scRNA-seq data indicated clear Epithelial, Endothelial and Mesenchymal clusters. **A)** Marker genes were used to initially assign the initial clustering into epithelial, endothelial, and mesenchymal clusters for downstream analysis. Marker gene expression by cluster is displayed in a heatmap, where higher expression is represented as a darker color. The known cell type that corresponds to the marker gene is indicated above the brackets at the top of the heatmap. **B)** UMAP embedding of all cells colored by developmental timepoint.

**Figure S2.**
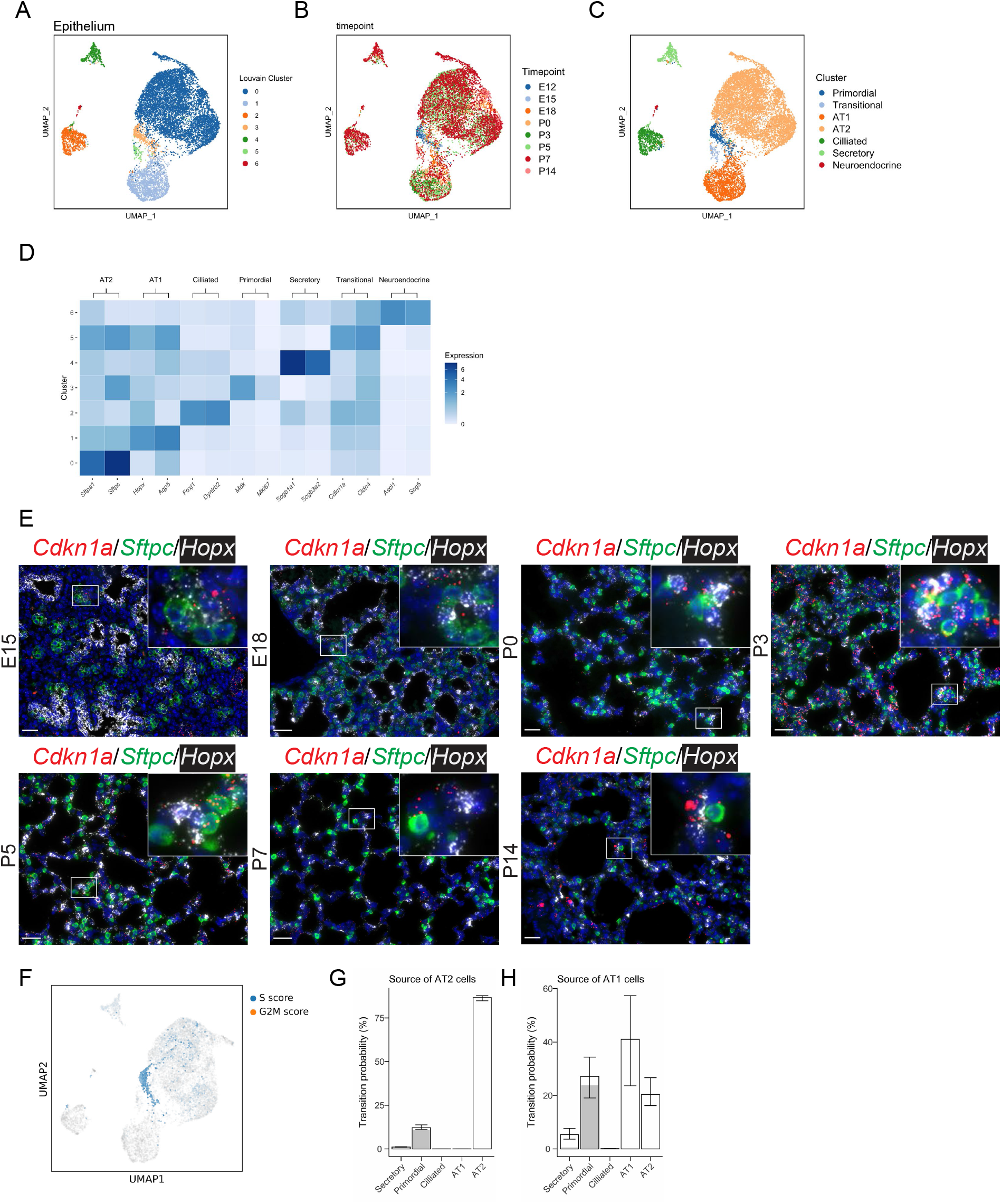
scRNAseq of the developing lung epithelium identifies the appearance of distinct cell types by E18. **A)** UMAP embedding of 11,456 cells from the developing lung epithelium, colored by Louvain cluster. **B)** UMAP embedding colored by timepoint. **C)** UMAP embedding colored by assigned cell-type. **D)** Clusters were assigned to cell types based on known marker gene expression. Maker gene expression by cluster is displayed in a heatmap, where higher expression is represented as a darker color. The cell type that corresponds to the marker gene is indicated above the brackets at the top of the heatmap. **E)** RNA in situ hybridization was used to identify cells that are expressing *Cdkn1a* (red, transitional cell marker), *Sftpc* (green, AT2 cell marker), and *Hopx* (white, AT1 cell marker). Scale bar = 25 µm. **F)** Cell cycle score analysis (a multi-gene metric) was performed with scVelo and plotted on a UMAP embedding. The darker the color, the greater the S or G2M cell cycle score. **G-H)** Cell fate probability was calculated with CellRank based on RNA expression profiles and RNA velocity to predict the source of **G)** AT1 cells and **H)** AT2 cells.

**Figure S3.**
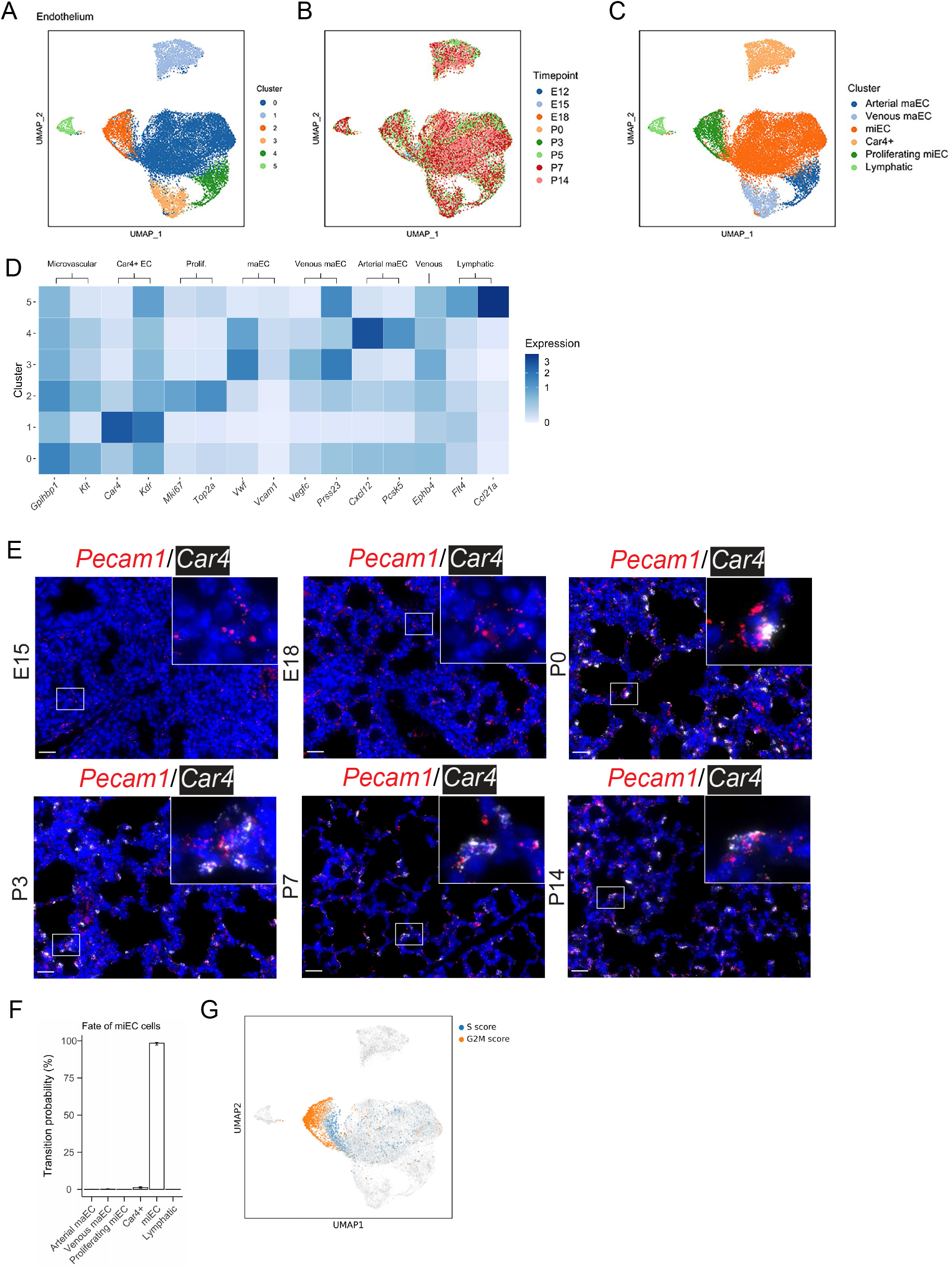
scRNA seq on the developing lung endothelium identifies early population stability, and an appearance of *Car4*+ cells at E18. **A)** UMAP embedding of 22,237 cells from the developing lung endothelium, colored by Louvain cluster. **B)** UMAP embedding colored by timepoint. **C)** UMAP embedding colored by identified cell-type. **D)** Clusters were assigned to cell types based on known marker gene expression. Marker gene expression by cluster is displayed in a heatmap, where higher expression is represented as a darker color. The cell type that corresponds to the marker gene is indicated above the brackets at the top of the heatmap. **E)** RNA in situ hybridization was used to identify cells that are expressing *Pecam1* (red, endothelial cell marker) and *Car4* (white, *Car4*+ cell marker). Scale bar = 25 µm. **F)** Cell fate probability was calculated with CellRank based on RNA expression profiles and RNA velocity to predict the fate of the microvascular endothelial cells (miEC). **G)** Cell cycle score analysis (a multi-gene metric) was performed with scVelo and plotted on a UMAP embedding. The darker the color, the greater the S or G2M cell cycle score.

**Figure S4.**
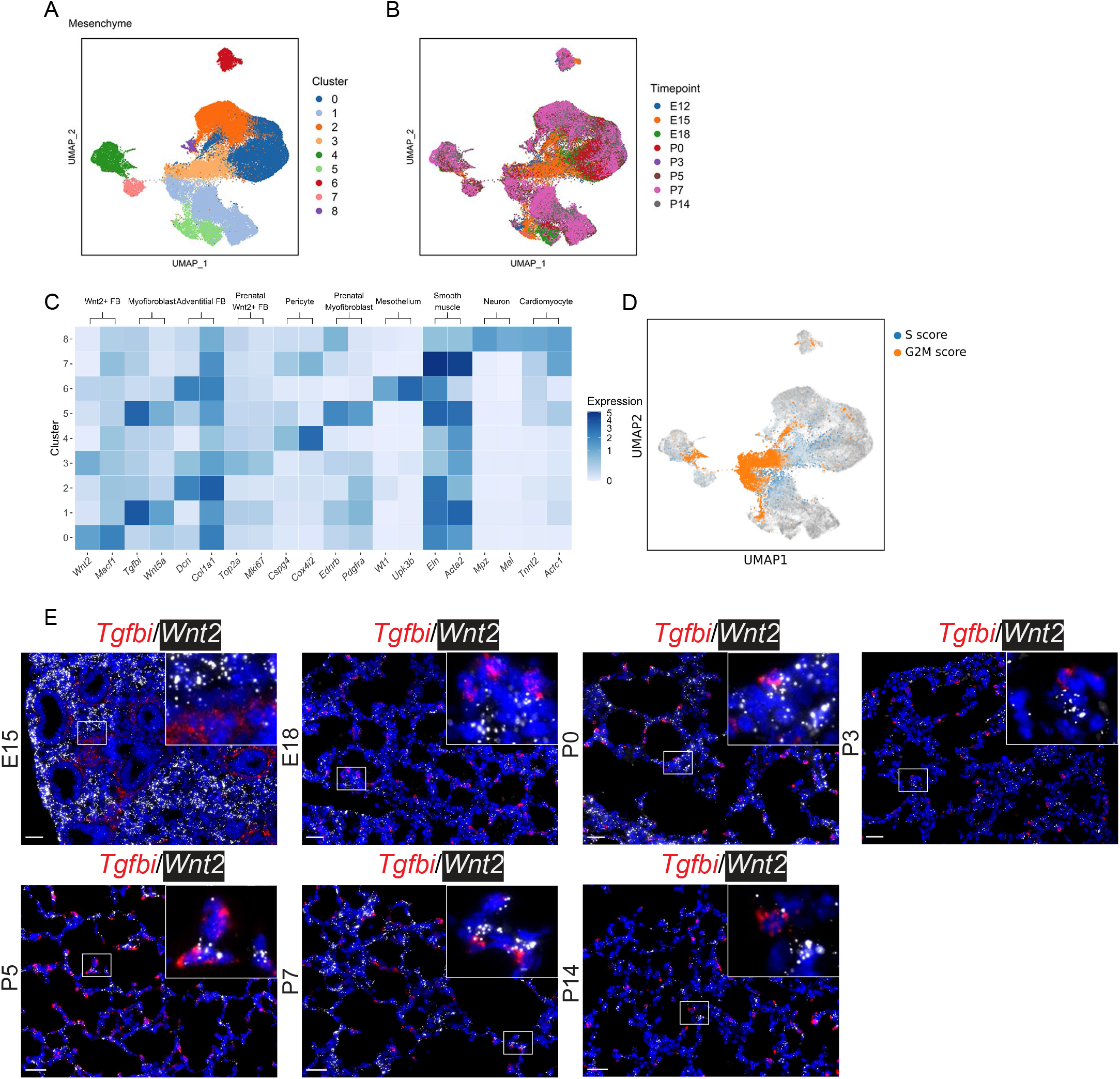
scRNA seq on the developing lung mesenchyme identifies broad changes over developmental time. **A)** UMAP embedding of 58,545 cells from the developing lung mesenchyme, colored by Louvain cluster. **B)** UMAP embedding colored by timepoint. **C)** Clusters were assigned to cell types based on known marker gene expression. Marker gene expression by cluster is displayed in a heatmap, where higher expression is represented as a darker color. The cell type that corresponds to the marker gene is indicated above the brackets at the top of the heatmap. **D)** Cell cycle score analysis (a multi-gene metric) was performed with scVelo and plotted on a UMAP embedding. The darker the color, the greater the S or G2M cell cycle score. **E)** RNA in situ hybridization (ISH) was used to identify cells that are expressing *Tgfbi* (red, myofibroblast marker) and *Wnt2* (white, *Car4*+ cell marker). Scale bar = 25 µm.

**Figure S5.**
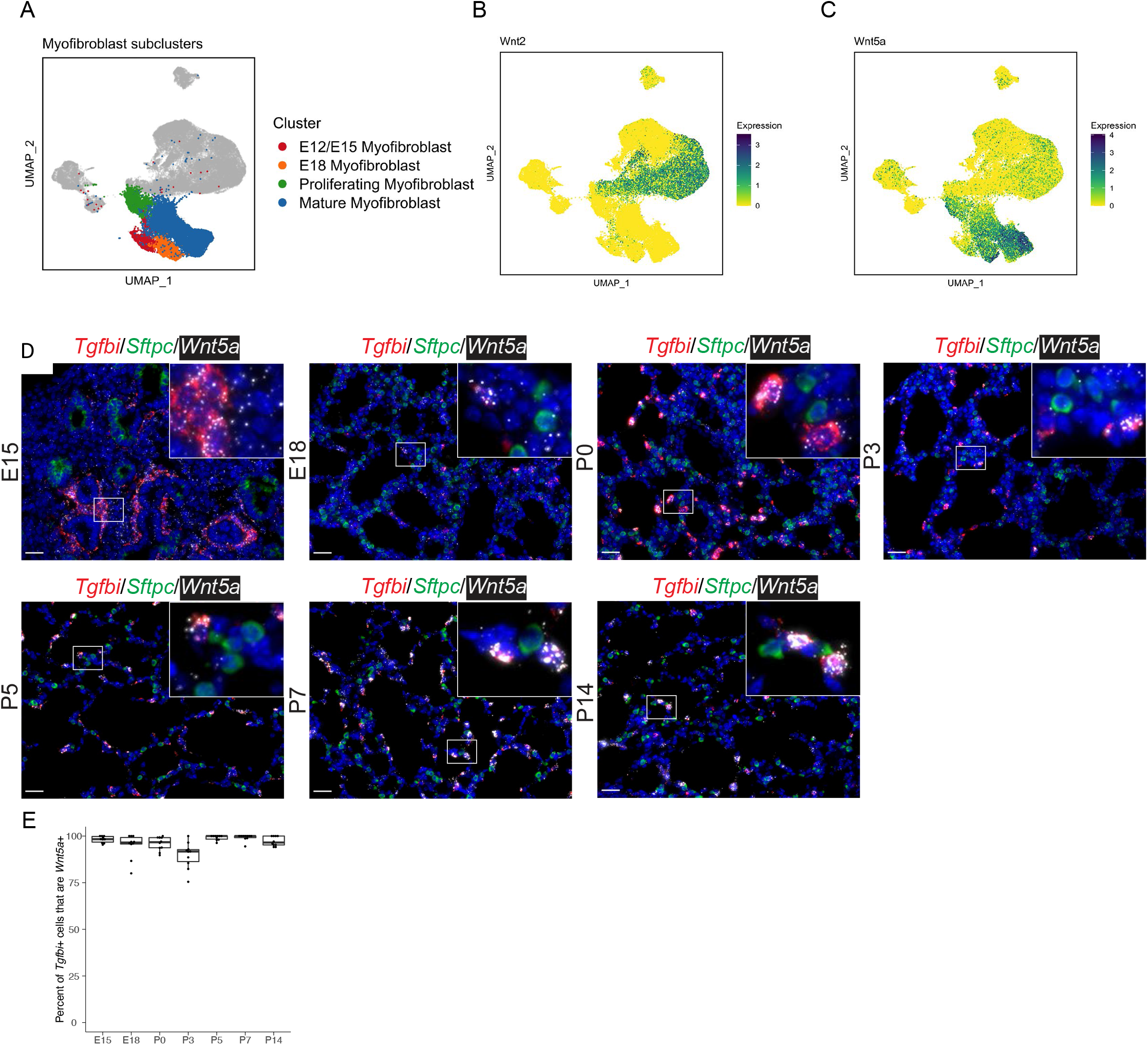
*Tgfbi+* myofibroblasts express *Wnt5a* after E15. **A)** UMAP embedding of the single-cell RNA Seq from the developing lung mesenchyme, with myofibroblast sub-clusters colored. **B)** UMAP embedding where individual cells are colored by the expression of *Wnt2* and **C)** *Wnt5a*. Darker colors indicate higher expression. **D)** RNA in situ hybridization (ISH) was used to identify cells that are expressing *Tgfbi* (red, myofibroblast marker), *Sftpc* (green, AT2 cell marker), and *Wnt5a* (white). Scale bar = 25 µm. **E)** Quantification of the ISH was performed using HALO and the percentage of *Tgfbi*+ cells that are *Wnt5a*+ was plotted. Boxplots represent the summary data and each dot represents the percentage of *Tgfbi*+*Wnt5a*+ / *Tgfbi*+ cells per image.

**Figure S6.**
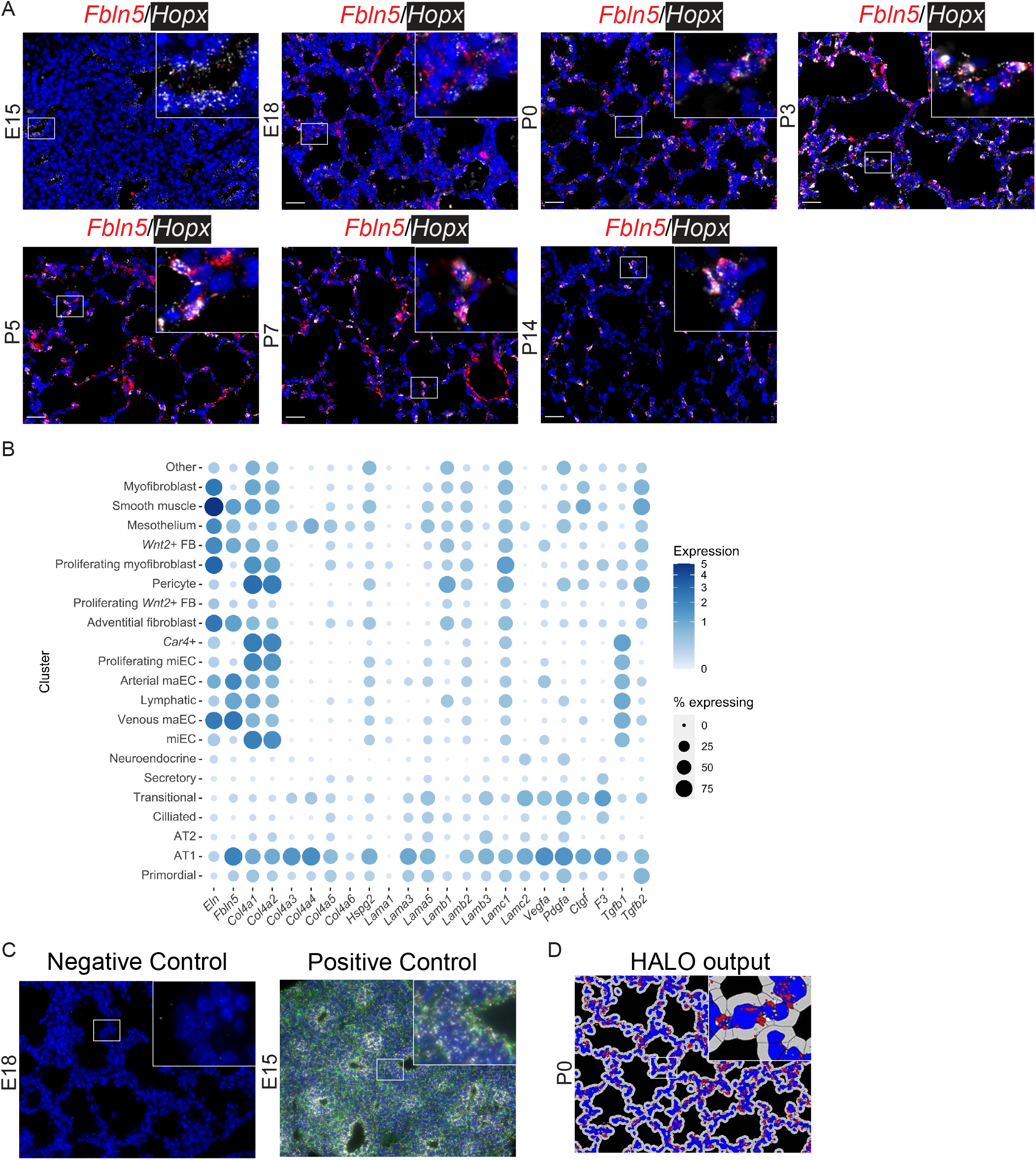
AT1 cells marked by *Hopx* express *Fbln5*. **A)** RNA in situ hybridization (ISH) was used to identify cells that are expressing *Fbln5* (red), and *Hopx* (white, AT1 cell marker). Scale bar = 25 µm. **B)** Heat map showing gene expression in every cluster of cells at all timepoints. The color intensity of the circles indicates the expression level and the size indicates the proportion of cells within a cluster that express a particular gene. **C)** All ISH experiments were accompanied with slides stained with positive and negative controls, a representative image of each is presented here. Both images were captured with identical exposure settings. **D)** HALO automated image analysis software (Indica Labs) was used for analysis of all ISH images. An example of HALO output for the P0 in panel A above is shown.

## References

Ahlmann-Eltze, C., and Huber, W. (2020). glmGamPoi: Fitting Gamma-Poisson Generalized Linear Models on Single Cell Count Data. Bioinformatics.

Barkauskas, C.E., Cronce, M.J., Rackley, C.R., Bowie, E.J., Keene, D.R., Stripp, B.R., Randell, S.H., Noble, P.W., and Hogan, B.L. (2013). Type 2 alveolar cells are stem cells in adult lung. J Clin Invest 123, 3025–3036.

Benjamin, J.T., Plosa, E., Sucre, J., van der Meer, R., Dave, S., Gutor, S.S., Nichols, D., Gulleman, P., Jetter, C., Han, W., et al. (2020). Neutrophilic inflammation during lung development disrupts elastin assembly and predisposes adult mice to COPD. J Clin Invest.

Benjamin, J.T., van der Meer, R., Im, A.M., Plosa, E.J., Zaynagetdinov, R., Burman, A., Havrilla, M.E., Gleaves, L.A., Polosukhin, V.V., Deutsch, G.H., et al. (2016). Epithelial-Derived Inflammation Disrupts Elastin Assembly and Alters Saccular Stage Lung Development. Am J Pathol 186, 1786–1800.

Bergen, V., Lange, M., Peidli, S., Wolf, F.A., and Theis, F.J. (2020). Generalizing RNA velocity to transient cell states through dynamical modeling. Nat Biotechnol 38, 1408–1414.

Bourbon, J., Boucherat, O., Chailley-Heu, B., and Delacourt, C. (2005). Control mechanisms of lung alveolar development and their disorders in bronchopulmonary dysplasia. Pediatr Res 57, 38R–46R.

Choi, J., Park, J.E., Tsagkogeorga, G., Yanagita, M., Koo, B.K., Han, N., and Lee, J.H. (2020). Inflammatory Signals Induce AT2 Cell-Derived Damage-Associated Transient Progenitors that Mediate Alveolar Regeneration. Cell Stem Cell 27, 366–382 e367.

Desai, T.J., Brownfield, D.G., and Krasnow, M.A. (2014). Alveolar progenitor and stem cells in lung development, renewal and cancer. Nature 507, 190–194.

Endale, M., Ahlfeld, S., Bao, E., Chen, X., Green, J., Bess, Z., Weirauch, M.T., Xu, Y., and Perl, A.K. (2017). Temporal, spatial, and phenotypical changes of PDGFRalpha expressing fibroblasts during late lung development. Dev Biol 425, 161–175.

Finak, G., McDavid, A., Yajima, M., Deng, J., Gersuk, V., Shalek, A.K., Slichter, C.K., Miller, H.W., McElrath, M.J., Prlic, M., et al. (2015). MAST: a flexible statistical framework for assessing transcriptional changes and characterizing heterogeneity in single-cell RNA sequencing data. Genome Biol 16, 278.

Frank, D.B., Penkala, I.J., Zepp, J.A., Sivakumar, A., Linares-Saldana, R., Zacharias, W.J., Stolz, K.G., Pankin, J., Lu, M., Wang, Q., et al. (2019). Early lineage specification defines alveolar epithelial ontogeny in the murine lung. Proc Natl Acad Sci U S A 116, 4362–4371.

Gillich, A., Zhang, F., Farmer, C.G., Travaglini, K.J., Tan, S.Y., Gu, M., Zhou, B., Feinstein, J.A., Krasnow, M.A., and Metzger, R.J. (2020). Capillary cell-type specialization in the alveolus. Nature 586, 785–789.

Goss, K. (2018). Long-term pulmonary vascular consequences of perinatal insults. J Physiol.

Hafemeister, C., and Satija, R. (2019). Normalization and variance stabilization of single-cell RNA-seq data using regularized negative binomial regression. Genome Biol 20, 296.

Hogan, B. (2018). Stemming Lung Disease? N Engl J Med 378, 2439–2440.

Kawasaki, T., Nishiwaki, T., Sekine, A., Nishimura, R., Suda, R., Urushibara, T., Suzuki, T., Takayanagi, S., Terada, J., Sakao, S., et al. (2015). Vascular Repair by Tissue-Resident Endothelial Progenitor Cells in Endotoxin-Induced Lung Injury. Am J Respir Cell Mol Biol 53, 500–512.

Kimmel, J.C., Penland, L., Rubinstein, N.D., Hendrickson, D.G., Kelley, D.R., and Rosenthal, A.Z. (2019). Murine single-cell RNA-seq reveals cell-identity- and tissue-specific trajectories of aging. Genome Res 29, 2088–2103.

Kobayashi, Y., Tata, A., Konkimalla, A., Katsura, H., Lee, R.F., Ou, J., Banovich, N.E., Kropski, J.A., and Tata, P.R. (2020). Persistence of a regeneration-associated, transitional alveolar epithelial cell state in pulmonary fibrosis. Nat Cell Biol 22, 934–946.

La Manno, G., Soldatov, R., Zeisel, A., Braun, E., Hochgerner, H., Petukhov, V., Lidschreiber, K., Kastriti, M.E., Lonnerberg, P., Furlan, A., et al. (2018). RNA velocity of single cells. Nature 560, 494–498.

Lange M B.V., Klein M, Setty M, Reuter B, Bakhti M, Lickert H, Ansari M, Schniering J, Schiller HB, Pe’er D, Theis FJ (2020). CellRank for directed single-cell fate mapping. bioRxiv 2020.10.19.345983.

Lee, J.H., Tammela, T., Hofree, M., Choi, J., Marjanovic, N.D., Han, S., Canner, D., Wu, K., Paschini, M., Bhang, D.H., et al. (2017). Anatomically and Functionally Distinct Lung Mesenchymal Populations Marked by Lgr5 and Lgr6. Cell 170, 1149–1163 e1112.

McInnes L H.J., Melville J. (2018). UMAP: Uniform Manifold Approximation and Projection for Dimension Reduction. arXiv.

Miner, J.H., Lewis, R.M., and Sanes, J.R. (1995). Molecular cloning of a novel laminin chain, alpha 5, and widespread expression in adult mouse tissues. J Biol Chem 270, 28523–28526.

Miner, J.H., Patton, B.L., Lentz, S.I., Gilbert, D.J., Snider, W.D., Jenkins, N.A., Copeland, N.G., and Sanes, J.R. (1997). The laminin alpha chains: expression, developmental transitions, and chromosomal locations of alpha1-5, identification of heterotrimeric laminins 8-11, and cloning of a novel alpha3 isoform. J Cell Biol 137, 685–701.

Morrisey, E.E., and Hogan, B.L. (2010). Preparing for the first breath: genetic and cellular mechanisms in lung development. Dev Cell 18, 8–23.

Nguyen, N.M., Kelley, D.G., Schlueter, J.A., Meyer, M.J., Senior, R.M., and Miner, J.H. (2005). Epithelial laminin alpha5 is necessary for distal epithelial cell maturation, VEGF production, and alveolization in the developing murine lung. Dev Biol 282, 111–125.

Nguyen, N.M., and Senior, R.M. (2006). Laminin isoforms and lung development: all isoforms are not equal. Dev Biol 294, 271–279.

Niethamer, T.K., Stabler, C.T., Leach, J.P., Zepp, J.A., Morley, M.P., Babu, A., Zhou, S., and Morrisey, E.E. (2020). Defining the role of pulmonary endothelial cell heterogeneity in the response to acute lung injury. Elife 9.

Plosa, E.J., Benjamin, J.T., Sucre, J.M., Gulleman, P.M., Gleaves, L.A., Han, W., Kook, S., Polosukhin, V.V., Haake, S.M., Guttentag, S.H., et al. (2020). beta1 Integrin regulates adult lung alveolar epithelial cell inflammation. JCI Insight 5.

Schuger, L., O’Shea, S., Rheinheimer, J., and Varani, J. (1990). Laminin in lung development: effects of anti-laminin antibody in murine lung morphogenesis. Dev Biol 137, 26–32.

Schuler, B.A., Habermann, A.C., Plosa, E.J., Taylor, C.J., Jetter, C., Negretti, N.M., Kapp, M.E., Benjamin, J.T., Gulleman, P., Nichols, D.S., et al. (2020). Age-determined expression of priming protease TMPRSS2 and localization of SARS-CoV-2 in lung epithelium. J Clin Invest.

Stouch, A.N., McCoy, A.M., Greer, R.M., Lakhdari, O., Yull, F.E., Blackwell, T.S., Hoffman, H.M., and Prince, L.S. (2016). IL-1beta and Inflammasome Activity Link Inflammation to Abnormal Fetal Airway Development. J Immunol 196, 3411–3420.

Stuart, T., Butler, A., Hoffman, P., Hafemeister, C., Papalexi, E., Mauck, W.M., 3rd, Hao, Y., Stoeckius, M., Smibert, P., and Satija, R. (2019). Comprehensive Integration of Single-Cell Data. Cell 177, 1888–1902 e1821.

Sucre, J.M.S., Vickers, K.C., Benjamin, J.T., Plosa, E.J., Jetter, C.S., Cutrone, A., Ransom, M., Anderson, Z., Sheng, Q., Fensterheim, B.A., et al. (2020). Hyperoxia Injury in the Developing Lung is Mediated by Mesenchymal Expression of Wnt5A. Am J Respir Crit Care Med.

Vila Ellis, L., Cain, M.P., Hutchison, V., Flodby, P., Crandall, E.D., Borok, Z., Zhou, B., Ostrin, E.J., Wythe, J.D., and Chen, J. (2020). Epithelial Vegfa Specifies a Distinct Endothelial Population in the Mouse Lung. Dev Cell 52, 617–630 e616.

Warburton, D., El-Hashash, A., Carraro, G., Tiozzo, C., Sala, F., Rogers, O., De Langhe, S., Kemp, P.J., Riccardi, D., Torday, J., et al. (2010). Lung organogenesis. Curr Top Dev Biol 90, 73–158.

Young, M.D., and Behjati, S. (2020). SoupX removes ambient RNA contamination from droplet-based single-cell RNA sequencing data. Gigascience 9.

Zacharias, W.J., Frank, D.B., Zepp, J.A., Morley, M.P., Alkhaleel, F.A., Kong, J., Zhou, S., Cantu, E., and Morrisey, E.E. (2018). Regeneration of the lung alveolus by an evolutionarily conserved epithelial progenitor. Nature 555, 251–255.

Zepp, J.A., Zacharias, W.J., Frank, D.B., Cavanaugh, C.A., Zhou, S., Morley, M.P., and Morrisey, E.E. (2017). Distinct Mesenchymal Lineages and Niches Promote Epithelial Self-Renewal and Myofibrogenesis in the Lung. Cell 170, 1134–1148 e1110.

